# The P2RX7B splice variant modulates osteosarcoma cell behaviour and metastatic properties

**DOI:** 10.1101/2021.05.07.443092

**Authors:** Luke Tattersall, Karan M. Shah, Darren L. Lath, Archana Singh, Jennifer M. Down, Elena De Marchi, Alex Williamson, Francesco Di Virgilio, Dominique Heymann, Elena Adinolfi, William D. Fraser, Darrell Green, Michelle A. Lawson, Alison Gartland

**Author notes:** Twitter: @DrLTattersall24, @k_m_shah @purineslab @lawson_ma, @ProfAllieG.

## Abstract

Osteosarcoma (OS) is the most common type of primary bone cancer affecting children and adolescents. OS has a high propensity to spread, meaning the disease is often incurable and fatal. There have been no improvements in survival rates for decades. This highlights an urgent need for development of novel therapeutic strategies. In this study, we have produced *in vitro* and *in vivo* data that demonstrates the role of purinergic signalling, specifically, the B isoform of the purinergic receptor P2RX7 (herein termed “ P2RX7B”), in OS progression and metastasis. Our data shows that P2RX7B expression confers a survival advantage in TE85+P2RX7B and MNNG-HOS+ P2RX7B human OS cell lines *in vitro* that is minimised following treatment with A740003, a specific P2RX7 antagonist. P2RX7B expression reduced cell adhesion and P2RX7B activation promoted invasion and migration *in vitro*, suggesting a probable metastatic phenotype. Using an *in vivo* OS xenograft model, MNNG-HOS+P2RX7B tumours exhibited ectopic bone formation that was abrogated with A740003 treatment. An increased metastatic phenotype was further demonstrated *in vivo* as expression of P2RX7B in primary tumour cells increased the propensity of the tumour to metastasise to the lungs. RNA-seq identified a novel gene axis, *FN1*/*LOX*/*PDGFB*/*IGFBP3*/*BMP4*, downregulated in response to A740003 treatment. In conclusion, our data indicates for the first time a role for P2RX7B in OS tumour growth, progression and metastasis. We show that P2RX7B is a potential therapeutic target in human OS.

**Novelty and Impact:** We provide evidence for the pro-tumorigenic role of the B isoform of the P2RX7 purinergic receptor in osteosarcoma (OS). In addition to increasing proliferation, P2RX7B increases the cancerous properties of OS cells, reducing adhesion and increasing migration and invasion. *In vivo*, P2RX7B does not affect primary tumour growth, but does lead to an increased propensity to metastasize. RNA-seq revealed a new axis of oncogenic genes inhibited by the P2RX7 antagonist and this data could potentially lead to new targets for OS treatment.

## Introduction

OS is the most common type of primary bone cancer affecting children and adolescents, it is a rare and often fatal disease^1^. OS is highly resistant to radiotherapy, therefore treatment involves complex chemotherapy and surgery, often amputation. Survival statistics have remained static with no real advances in treatment options for decades. The 5-year overall survival rate has remained static for decades, at 53% for patients under 40 years of age^2^, however, with metastasis this figure decreases to around 20%^3^. These statistics highlight an urgent need for development of novel therapeutic strategies for OS treatment.

Purinergic receptors are expressed and play a role in the progression of several cancers^4^. Several studies have identified adenosine triphosphate (ATP) to be at a high concentration in the tumour microenvironment^5, 6^, yet undetectable in surrounding healthy tissue^7^. These findings are particularly pertinent in the case of the bone tumour microenvironment where purinergic receptors are known to be expressed by bone cells that play a key role in normal bone physiology^8, 9^ and where mechanical loading can stimulate ATP release from bone cells^9-11^. ATP when broken down to adenosine can act as an immunosuppressant and chemotherapy treatments can cause the release of ATP from dying cells^12^. This release can modify the tumour microenvironment in favour of cancer growth and potentially result in disease progression. Purinergic receptor P2RX7 is an ATP gated ion channel and has received an increasing amount of attention due to both *in vitro* and *in vivo* evidence of its overexpression and function in carcinogenesis of a range of different tumours, including bone sarcomas ^4, 12-15^. P2RX7 contributes to various oncogene-like properties such as enhancing tumour cell growth and metabolic activity, increasing invasiveness and metastatic potential by acting in an autocrine/paracrine manner on surrounding cells and by acting as a stimulant for the release of angiogenic factors^14^. Nine different naturally occurring human P2RX7 splice variants (P2RX7A to P2RX7J) have been identified each showing differing biological characteristics^16, 17^. P2RX7 isoform expression, phenotype and level of activation can determine the functional role that P2RX7 has in the tumour microenvironment. The full length P2RX7A variant has the unique ability to form a large transmembrane pore in response to high levels of ATP resulting in cell death. P2RX7B preserves an intron between exons 10 and 11 that causes the insertion of a stop codon that eliminates translation of the last 249 amino acids of the C terminus and the addition of an extra 18 amino acids after residue 346. These alterations still enable receptor stimulation by ATP/Benzoyl ATP (BzATP) and importantly efficacy of P2RX7 antagonists^17^. Of the splice variants, P2RX7B is unique in that it retains its ability to act as a functional ion channel^17, 18^ and shares a similar tissue distribution as the full length receptor^19^. Due to the truncated nature of the P2RX7B it naturally lacks the pore function with a “ non-functional” phenotype found to be essential for tumour cell growth^20^ and more recently other features including response to treatment, invasion and promotion of metastasis^21^. In lung adenocarcinoma patient samples, high P2RX7B expression correlated with reduced immune cell infiltration, suggesting it could be a potential biomarker^22^. Whilst in acute myeloid leukaemia, P2RX7B expression was increased in relapsing patients and downregulated in patients in remission, suggesting high levels of P2RX7B may contribute towards chemoresistance. This was proposed to be due to the absence of the full-length P2RX7B receptor pore, as when the full length receptor was present chemotherapy uptake was increased. The study further showed that treatment *in vivo* with combined chemotherapy and P2RX7 antagonist was an effective strategy acting on both isoforms, reducing tumour growth and isoform expression in the primary tumour ^23^.

Previous studies by our lab and others have shown a potential role for targeting P2RX7 as a therapeutic option in OS^15, 24^. When assessing different P2RX7 splice variants in TE85 human OS cells, P2RX7B provided a growth advantage, and when expressed in OS patient samples correlated with higher cell numbers suggesting OS phenotypes can be influenced by PR2X7B^15^. More recently it has been suggested that the full length P2RX7 could promote OS cell growth and metastasis both *in vitro* and *in vivo*^24^. In this study we have built upon both of these previous studies and explored the role of the specific P2RX7B splice variant in OS both *in vitro* and *in vivo*.

## Materials and methods

### Cell culture and transfections

The human OS cell lines HOS-TE85^25^ (referred to heroin as TE85) and MNNG-HOS^26^ were kindly provided from Professor Jim Gallagher (Liverpool, UK) and Professor Dominique Heymann (INSERM, France), respectively. HEK-293+P2RX7A cells were kindly provided from the laboratory of Dr Elena Adinolfi (Ferrara, Italy). All cell lines were authenticated using STR profiling during the project period/within the last three years. Cell lines were maintained in DMEM©+Glutamax™ (Life Technologies Paisley, UK) medium containing 10% FBS with 1% penicillin/streptomycin antibiotics (Life Technologies Paisley, UK) (referred to as complete medium). TE85 cells previously stably transfected with P2RX7B^15^ were maintained in standard culture medium plus 0.2 mg/ml hygromycin. MNNG-HOS cells containing a plasma membrane luciferase (PmeLUC) from the lab of Professor Francesco Di Virgilio^27^ were further transfected with the mammalian expression vector pcDNA3 containing P2RX7B^15^ using Lipofectamine and maintained under 0.8 mg/ml G418 selection for PmeLUC and 0.2 mg/ml hygromycin for P2RX7B. All cells were grown in a 37°C in a humidified 5% CO_2_ incubator and were regularly mycoplasma tested to ensure that and all experiments were performed with mycoplasma-free cells.

### Fluo-4 Direct™ Calcium Assay

Changes in intracellular calcium concentrations were measured using Fluo-4 Direct™ Calcium Assay (ThermoFisher Scientific, UK) in accordance with the manufacturer’s instructions. In brief, 15,000 cells were loaded with Fluo-4 Direct™ calcium reagent, left to incubate for 1 hour at 37°C before being stimulated with 100 µM BzATP (Sigma, UK) and then 0.8 µM ionomycin (Sigma, UK). Excitation ratio and emission wavelength were 490 nm and 525 nm respectively.

### Measurement of plasma membrane permeabilization

Increases in plasma membrane permeability were measured by monitoring ethidium bromide up-take. Briefly,15,000 cells were incubated at 37°C with or without 10 µM of the A740003 (Tocris, UK) for 1 hour in Hank’s Balanced Salt Solution (HBSS) (ThermoFisher Scientific). Cells were then stimulated with 300 µM BzATP diluted in ultra-pure ethidium bromide to a final concentration of 100 µM. Induction of P2RX7 pore formation was then detected at 360 nm excitation and 580 nm emission for 45 minutes after an initial 5-minute baseline reading, with readings taken every 2 minutes.

### Proliferation assay

The effect of P2RX7B expression and inhibition on OS cell proliferation was assessed using the CellTiter 96® AQueous One Solution Cell Proliferation Assay (Promega, UK), TE85, TE85+P2RX7B, MNNG-HOS and MNNG-HOS+P2RX7B cell lines were seeded at 5,000 and 2,500 cells per well, respectively, in a 96-well plate in 100 μL phenol-free DMEM (Life Technologies) complete medium. After 24 hours, cells were washed twice with PBS and the medium was changed to medium containing 0.5% FBS. At each time-point on days 0, 1, 3, 5 and 7, MTS was diluted 1:5 in phenol-free DMEM with 100 μL added directly making a total volume of 200 μL. The cells were incubated for 3 hours at 37°C. The absorbance was read at 490 nm using the SpectraMax M5e Microplate Reader (Molecular Devices). For inhibition studies the same protocol was performed with 100 μM A740004 added after 24 hours with absorbance read at day 3.

### Cell adhesion assay

96-well plates coated with 50µg/mL Type 1 rat tail collagen (Sigma, UK) were seeded with 7,500 cells per well in medium containing 0.5% FBS. After incubation for 4 hours at 37°C the wells were washed 4 times with PBS to removed unattached cells. Cells were then lysed using 50 µL lysis buffer (20 mM tris, 0.05 M MgCl_2_) and Quant-iT™ PicoGreen® dsDNA Reagent (ThermoFisher Scientific) was then added to detect the DNA of the lysed remaining cells. Fluorescence was detected and quantified at 485 nm excitation and 530 nm emission.

### Cell migration (scratch assay)

12-well plates were seeded with 200,000 cells in complete medium and left overnight to form a confluent monolayer. The medium was then changed to complete medium containing 5 µg/mL mitomycin C and cells incubated for 2 hours at 37°C. A scratch was then made using a 10μL pipette tip down the centre of the well. The wells were washed twice with PBS and fresh 0.5% FBS medium added. Images were then taken every 2 hours for a 24-hour period. For studies activating the P2RX7B 10 µM BzATP was used again in low 0.5% FBS medium. Scratch assay images were analysed using the automated T-scratch software^28^.

### Cell invasion

Matrigel (1.5 mg/mL) was added to Corning® FluoroBlok inserts and left to set for 2 hours at 37°C. Cells were incubated with 5 µg/mL mitomycin C for 2 hours before 100,000 cells were added to Matrigel-coated insert in serum free medium +/- 10 µM BzATP. 10% FBS was used as a chemoattractant in the lower chamber. After 24 hours the transwell inserts were washed twice in PBS and stained with 5 µM Calcein AM cell permeant dye for 30 minutes at 37°C. The inserts were then washed twice again in PBS, imaged and analysed using Image J^29^.

### RNA extraction and End Point PCR

RNA was extracted from cells using the ReliaPrep™ RNA Miniprep System kit in accordance with the manufacturer’s protocol (Promega). RNA was reverse transcribed using the Applied Biosystems™ high capacity RNA to cDNA™ Kit in accordance with the manufacturer’s protocol. End point PCR primers were designed using Pubmed gene sequences. P2RX7 mRNA expression was determined with two sets of primers; the first primer set was designed in the N terminal gene sequence with the forward primer on the exon boundary between exon 3 and 4 and the reverse between exon 7 and 8. This region is present on both the full length and truncated P2RX7B (Forward: 5’-TTGTGTCCCGAGTATCCCAC-3’ Reverse: 5’-TCAATGCCCATTATTCCGCC-3’ product length 413 bp,). The second primer set was designed further along the gene sequence with the both the forward primer and reverse primer designed to bind to exon 13 (Forward 5’-ACCAGAGGAGATACAGCTGC-3’ Reverse: 5’-TACTGCCCTTCACTCTTCGG-3’, product length 399 bp). This C-terminal region is only present on the full length P2RX7A and thereby not detected in the cells with the truncated P2RX7B. cDNA was amplified using Promega GoTaq Flexi DNA polymerase kit with primers (0.5 µM) and template cDNA (1 µg) added. The PCR samples were denatured for 2 minutes at 95°C for one cycle, followed by 35 cycles of denaturation at 90°C for 30 seconds, primer annealing temperature for 30 seconds, and extension at 72°C for 30 seconds. A final extension was then performed at 72°C for 5 minutes for one cycle before samples were held at 4°C.

### Quantitative RT-PCR (qRT-PCR)

qRT-PCR was performed using Taqman probes (Human P2RX7 Taqman® gene expression assay, ID: Hs00951600_m1 catalogue: 4351372, human HPRT Taqman® gene expression assay, ID: Hs02800695 catalogue: 1621448) in accordance with the manufacturer’s instructions on a Applied Biosystems 7900HT Real-Time PCR machine (Applied Biosystems™).

### RNA-seq of P2RX7B agonist and antagonist treated MNNG-HOS and MNNG-HOS+P2RX7B OS cell lines

MNNG-HOS and MNNG-HOS+P2RX7B cells were plated in a 6 well plate at a density of 500,000 cells and left for 24 hours to adhere. Cells were then treated with 10 µM BzATP, 10 µM A740003 or both BzATP and A740003 together (A740003 was pre incubated for 1 h before adding BzATP) 24 hours later RNA was extracted using the miRNeasy mini kit (Qiagen) according to the manufacturer’s protocol. RNA concentration and integrity were assessed on a NanoDrop 8000 Spectrophotometer (Thermo Fisher Scientific) and RNA was stored at -80 °C. The NEBNext ultra II RNA library prep kit (New England Biolabs) was used to generate poly(A)+ libraries. Sequencing was performed on a HiSeq 2500 (Illumina) using a 150 bp paired end (PE) metric.

### Bioinformatics

Fastq files were converted to fasta. Trim Galore was used to remove adapter sequences and reads <20 nucleotide. Trimmed reads were aligned to the human genome (v38) using HISAT2^30^. Transcripts were download from GENCODE (v28) and Ensembl (v92). Count matrices for transcripts were created using Kallisto^31^. Differentially expressed (DE) transcripts were determined using the DESeq2^32^ package in R (v1.2.10). DE mRNA according to log_2_ fold change ≥2, p ≤ 0.05 plus false discovery rate (FDR) <5% were selected for downstream analysis. Pathway analysis was used to determine the biological pathways of the different genes using the KEGG database.

### *In silico* analysis of OS datasets for P2RX7 expression

A retrospective analysis across publicly available datasets was performed to identify P2RX7 expression in human OS cell lines. Searches were performed in the Cancer Dependency Map using the 20Q4 depmap gene expression dataset (Broad Institute, https://depmap.org/portal/), R2 Genomics Analysis and Visualization Platform using datasets (GSE86109, GSE42352, GSE11414, GSE124768 (http://r2.amc.nl) and Ordino a visual cancer analysis database https://ordino.caleydoapp.org/ where the gene P2RX7 was selected and filtered for OS, the identified cell lines were then selected for analysis.

### *In vivo* animal studies

All animal experiments were performed using 7-9-week-old female BALB/c nude mice (Charles River Margate, UK) that were acclimatised for at least one week prior to experimental manipulation. Mice were housed in the same environmentally controlled conditions with a 12hr light/dark cycle at 22°C and free to access 2018 Teklad Global 18% Protein Rodent Diet containing 1.01% Calcium (Harlan Laboratories, UK) and water ad libitum in RB-3 cages. All procedures complied with the UK Animals (Scientific Procedures) Act 1986 and were reviewed and approved by the local Research Ethics Committee of The University of Sheffield (Sheffield, UK). Mice were anesthetized by inhalation of isoflurane and oxygen before injection of 250,000 MNNG-HOS or MNNG-HOS+P2RX7B cells onto the tibial surface. For inhibition studies, mice were randomly allocated into treatment groups (N=11-13 mice/group) 2 days after OS cell injection and received either vehicle (PBS+DMSO), or A740003 (50 µg/kg) by IP injection 3 times a week for up to 3 weeks OS tumours were measured by callipers at the end of the experiment prior to euthanasia, the legs and lungs were then harvested for analysis. For OS metastasis studies mice were anesthetized as above before 1.5 million MNNG-HOS or MNNG-HOS+P2RX7B cells were injected into the tail vein. After 2 days the mice were divided into 4 groups (N=6 mice/group) and treated with either vehicle (PBS+DMSO) or A740003 (50 µg/kg) by IP injection 3 times a week for 3 weeks, after euthanasia the lungs were harvested.

### Micro-CT analysis

Fixed tibiae were scanned using a SkyScan 1172 desktop μCT machine (Bruker) at a resolution of 8 μm with the X-ray source operating at 50 kV, 200μA and using a 0.5 mm aluminium filter and images were captured every 0.7°. Scanned images were reconstructed using Skyscan NRecon software (v. 1.6.9, Bruker, Belgium). The region of interest (ROI) for the total bone volume was selected to include both the tibia and fibula and was determined at the top of the bone as soon as the tibia enters the image to the lower point where the tibia and fibula meet.

### Histological studies on the primary tumour

Bones were fixed in neutral buffered formalin for 48 hours after which time they were transferred to 70% ethanol. Bones were then decalcified in 10% EDTA, embedded in paraffin and 5 μm sections produced. Sections were then dewaxed in xylene, rehydrated through graded alcohols before performing heat mediated antigen retrieval using a water bath at 80°C for 20 min with citrate buffer at pH6 (Abcam, Cambridge, UK). Endogenous peroxidase was blocked with 3% hydrogen peroxide (VWR, Lutterworth, UK) for 20 min at room temperature, washed twice in PBS-tween (PBST) and blocked in 1% Normal goat serum in PBST (Vector Laboratories, Peterborough, UK) for 20 minutes at room temperature. Primary rabbit anti-human Ki-67 antibody (Abcam 1 mg/mL) was added to the sections at a dilution of 1:100 in 1% casein. Sections were incubated for 1 hour at room temperature. After 2 washes in TBST, the sections were treated with biotinylated goat anti-rabbit IgG secondary antibody (Vector Laboratories) at 1:200 in 1% casein for 20 minutes at room temperature. Sections were washed twice in PBST then treated with an ABC kit (Vector Laboratories) for 20 min at room temperature and the bound antibody detected with Impact-DAB substrate-chromagen system (Vector Laboratories) for 5 min at room temperature. The sections were washed in tap water for 3 minutes, counter stained in Gills haematoxylin (VWR, Merck, Birmingham, UK) for 5 seconds, dehydrated through graded alcohols, and cleared with xylene. Coverslips were mounted using DPX. The slides were scanned using a Pannoramic 250 Flash III (3D HISTECH, Budapest, Hungary) and percentage of Ki-67 cells were quantified using QuPath software^33^. Tumours were further categorised histologically into high or low grade tumours using the median % Ki-67 value of (20.17%) as a cut off^34^.

### Histological studies for lung metastasis

Lungs were collected and fixed in 10% neutral buffered formalin for 48 hours before being changed into 70% ethanol and embedded in paraffin. To section the lungs 6 μm sections were cut every 100 μm deep to cover the entire lung. Sections were added to xylene twice for 5 minutes rehydrated through graded alcohols for 5 minutes each and tap water for 1 minute. The nuclei were then stained by Gill’s II haematoxylin for 90 seconds and washed using tap water for 3 minutes. 1% aqueous eosin (VWR, Merck, Birmingham, UK) with 1% calcium carbonate (Sigma, UK) used to stain the cytoplasm for 5 minutes. The slides were then quickly dehydrated through graded alcohols and cleared with xylene. Coverslips were mounted using DPX. Lung sections were visualised under a light microscope and were observed by two independent researchers blinded to the treatment groups for the presence of any OS metastatic nodules.

### Statistical analysis

All data was analysed using GraphPad Prism v7. Datasets were tested for normality prior to analysis and statistical significance was tested using parametric or non-parametric tests as appropriate for the specific dataset. For comparing two groups, either a paired or unpaired t-test was used depending upon the groups to be compared. Two-way ANOVA was used to determine the effect of P2RX7B expression on tumour-induced changes in bone parameters *in vivo*. Slopes were compared using linear regression for the wound closure migration assay and a Chi-squared was used for Ki-67 staining.

## Results

### P2RX7B expression and functional characteristics in OS cells

We have previously shown that the human OS cell line TE85 lacks endogenous P2RX7 protein expression^35^ and that overexpression of the P2RX7 in this cell line confers trophic activity, with the most efficient growth-promoting isoform being P2RX7B alone^15^. To further investigate the role of the P2RX7B in OS pathology both TE85 and the MNNG-HOS cell line (a chemically transformed cell line derived from the original TE85 cell line which, unlike TE85 cells, forms tumours in mice *in vivo*^36^) were utilised. Similar to TE85 cells, we did not detect endogenous P2RX7 expression in MNNG-HOS cells at the mRNA transcript level using end point and qRT-PCR (Figure 1A & 1B). We also used the Ca^2+^ influx and pore formation assays to test for the presence of functional P2RX7, the results of which support the lack of endogenous P2RX7 in these cells (Figure 1C & 1D). We therefore transfected both these cell lines with the truncated P2RX7B isoform (Figure 1A & 1B), and confirmed functional expression using P2RX7 Ca^2+^ signalling and pore formation (Figure 1C & 1D), the latter being unaltered as predicated. To further examine the expression of P2RX7 in OS cell lines, we performed retrospective *in silico* analysis using publicly available datasets. DepMap contained P2RX7 gene expression data for 14 OS cell lines, R2 genomics had data for 38 cell lines (across 4 datasets) and Ordino had data for 9 cell lines. The data was then arranged showing P2RX7 expression across the cell lines. From these datasets the expression data for cell lines relating to the HOS-TE85 family and its modified derivatives KHOS (viral), MNNG (chemical) and 143b (metastatic)^37, 38^ were consistently low for P2RX7 expression in comparison to other OS cell lines (Supplementary Figure 1A-E). Cell lines that were consistently high for P2RX7 expression were SJSA-1 (Supplementary Figure 1A-D), CAL72 (Supplementary Figure 1B & D) and MG63 (Supplementary Figure 1A-D & F).

**Figure 1:**
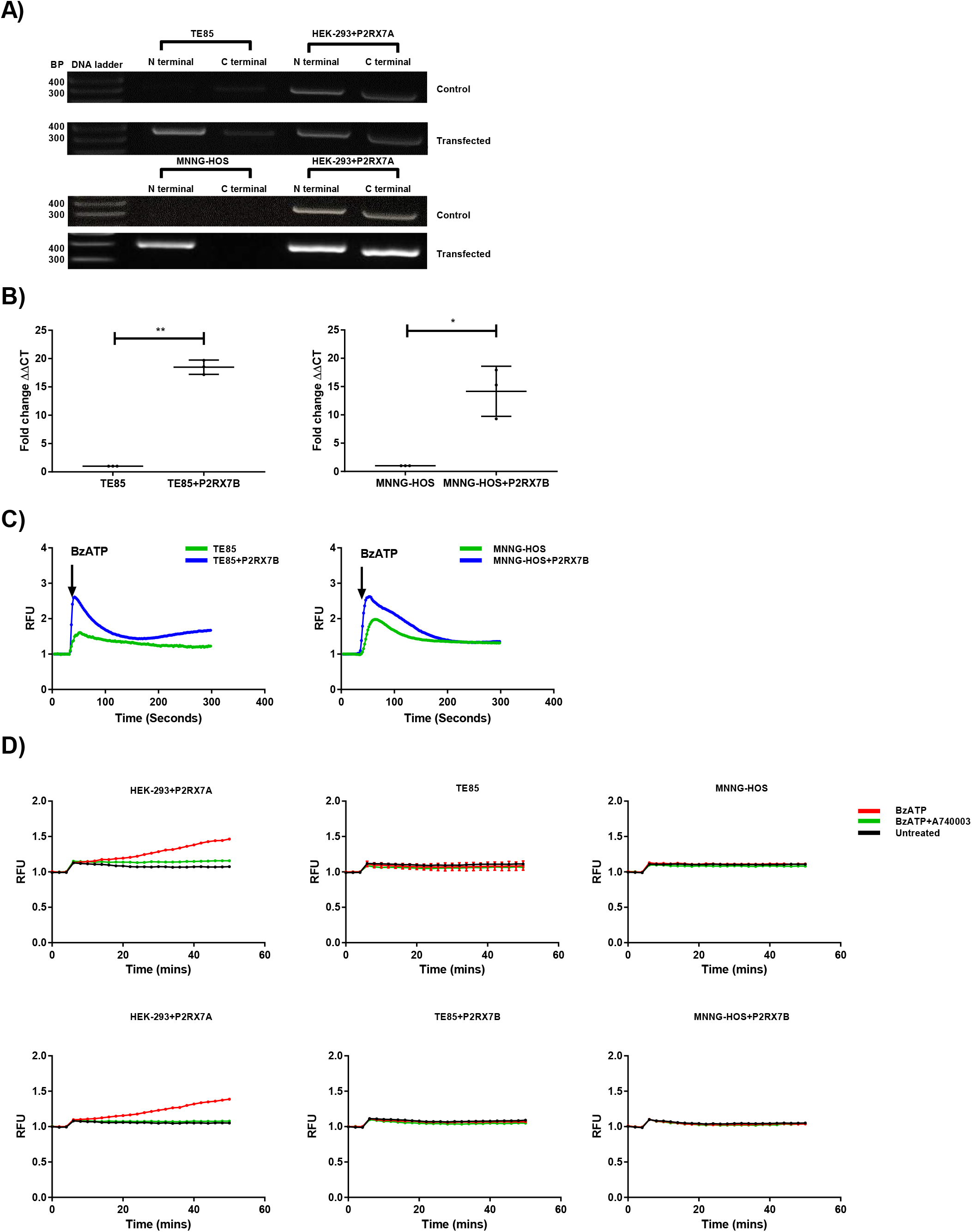
Characterising P2RX7B in TE85 and MNNG-HOS OS cells: **A)** Expression of P2RX7 mRNA in TE85, TE85+P2RX7B, MNNG-HOS and MNNG-HOS+P2RX7B cells with two different P2RX7 sets of primers binding to the P2RX7 N Terminal or C terminal, (product length 413 BP and 399 BP respectively) HEK-293+P2RX7A was used as a positive control. **B)** Expression of P2RX7 mRNA in TE85, TE85+P2RX7B, MNNG-HOS and MNNG-HOS+P2RX7B OS cells using qRT-PCR, HPRT was used as a housekeeping gene. **C)** Calcium assays were assessed in TE85, TE85+P2RX7B, MNNG-HOS and MNNG-HOS+P2RX7B cells by loading 15000 cells with Fluo-4 Direct™ calcium reagent, left to incubate for 1 hour at 37°C before been stimulated with 100 µM BzATP and then 0.8 µM ionomycin, the response was detected at excitation ratio and emission wavelength 490 nm and 525 nm respectively for five minutes. **D)** Measurement of plasma membrane permeabilization was performed on TE85, TE85+P2RX7B, MNNG-HOS and MNNG-HOS+P2RX7B cells using ethidium bromide uptake, 15000 cells in HBSS were incubated at 37°C with or without 10 μM of the P2RX7 inhibitor A740003 for 1 hour. BzATP was diluted in ultra-pure ethidium bromide to a final concentration of 300 μM BzATP and 100 μM ethidium bromide, the response was detected at 360 nm excitation and 580 nm emission for 45 minutes after an initial 5-minute baseline reading, with readings taken every 2 minutes, HEK-293+P2RX7A was used as a positive control. All data is from 3 biological repeats with 6 technical replicates per experiment and were compared using an unpaired T-test, * = P <0.05 ** = P<0.01.

### P2RX7B expression increases the growth and metastatic potential of OS cells

Transfection of P2RX7B into both OS cell lines caused an increase in proliferation under low serum conditions over a 7-day period. In TE85 cells transfected with P2RX7B (TE85+P2RX7B) there was a growth increase of 33% across day 3 (P=0.0001), 5 (P=0.0003) and 7 (P= 0.0011) (Figure 2Ai), The same increase in growth was also observed in MNNG-HOS cells transfected with P2RX7B (MNNG-HOS+P2RX7B) - 56% increase across day 3 (P=<0.0001), 5 (P=<0.0001), and 7 (P= <0.0001) (Figure 2Aiv). The P2RX7 antagonist A740003 reduced growth increase by 40% in TE85+P2RX7B (P= <0.0001, Figure 2Aii) and 52% in MNNG-HOS+P2RX7B cells (P= <0.0001, Figure 2Av). In both cell lines A740003 had no statistically significant effect on the parental cell lines not transfected with P2RX7B (Figure 2Aiii and 2Avi).

**Figure 2:**
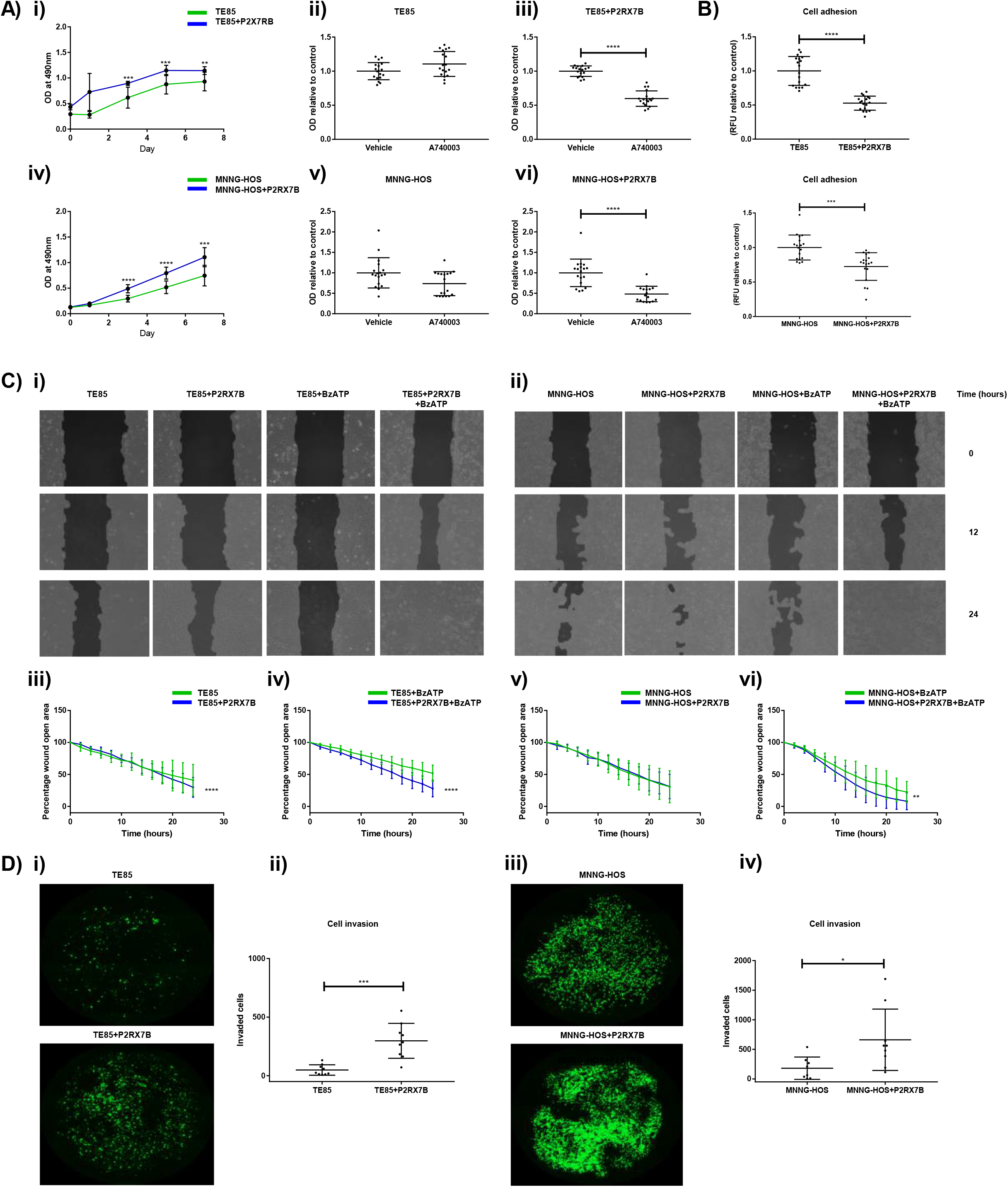
The effect P2RX7B expression on TE85 and MNNG-HOS OS cell proliferation, adhesion, migration and invasion. Cell proliferation was assessed using an MTS assay over 7 days with absorbance taken on day 0,1,3,57. For inhibition studies A740003 was added on day 1 and measured on day 3. **Ai)** TE85 compared to TE85+P2RX7B, **Aii)** TE85 treated with A740003, **Aiii)** TE85+P2RX7B treated with A740003 **Aiv)** MNNG-HOS compared to MNNG-HOS+P2RX7B **Av)** MNNG-HOS treated with A740003 **Avi)** MNNG-HOS+P2RX7B treated with A740003. Cell adhesion was assessed by plating 7500 cells into a 96 well plate pre-coated with type 1 rat tail collagen and left for 4 hours at 37°C, wells were washed 4 times with PBS and remaining attached cells lysed and detected using Quant-iT™ PicoGreen® dsDNA Reagent. Fluorescence was detected at excitation 485 nm and emission 530 nm. **B)** TE85 compared to TE85+P2RX7B and MNNG-HOS compared to MNNG-HOS+P2RX7B. Migration was assessed by plating 200,000 cells into a 12 well plate and left 24 hours to form a monolayer, the cells were incubated in 5 μg/ml mitomycin C for 2 hours and then scratched down the centre of the well using a 10 μL pipette tip, after washing twice with PBS the cells were left in medium containing 0.5% FBS with images taken every 2 hours for 24 hours All images were analysed, and pseudo coloured using Tscratch software **Ci)** Representative images for TE85 and TE85+P2RX7B individually and when stimulated with BzATP, with the corresponding wound closure data shown in **Ciii** and **CiV**. Representative images for MNNG-HOS and MNNG-HOS+P2RX7B individually and when stimulated with BzATP are shown in **Cii** with the corresponding wound closure data shown in **Cv** and **Cvi**. For invasion cells were incubated in culture with 5 μg/ml mitomycin C and left for 2 hours at 37°C. They were seeded into a 24 well plate at a density of 100,000 cells in serum free medium containing 10 μM BzATP, in an upper fluoroblok chamber pre-coated with 1.5 mg/ml matrigel. The medium in the lower chamber contained complete medium. After 24 hours the upper fluoroblok transwells were washed twice in PBS and left for 30 minutes in calcein AM cell permeant dye to stain live invaded cells. Images were taken covering 60% of the underside of the 24-well, this covered the entire surface of the smaller insert and were analysed using Image J. **Di)** Representative images of TE85 compared with TE85+P2RX7B with the number of invaded cells shown in **Dii**. Representative images of MNNG-HOS compared with MNNG-HOS+P2RX7B are shown in **Diii** with the number of invaded cells in **Div**. All data is from 3 biological repeats with 3-6 technical replicates per experiment and were compared using an unpaired T-test with migration slopes compared by linear regression * = P <0.05 ** = P<0.01 *** = P< 0.001 **** = P < 0.0001.

To assess the effect of P2RX7B expression on adhesion, type 1 collagen was used as an extracellular matrix (ECM) and the cells ability to adhere to it was measured. TE85+P2RX7B cells had a 47% reduction in cell adhesion (P=<0.0001, Figure Bi) and MNNG-HOS+P2RX7B cells had a 27% reduction in cell adhesion (P=<0.0001, Figure Bii) compared with their respective controls. To measure the effect of P2RX7B expression on OS cell migration a scratch assay was performed where the cells were monitored over a 24-hour period for “ wound” closure. After 24 hours under low serum conditions, TE85+P2RX7B showed a 12% increase in wound closure compared to TE85 cells and showed an overall difference when comparing the rate of closure (P=<0.0001, Figure 2Ci and 2Ciii). This difference was further increased to 24% when stimulated with 10 µM BzATP (P=<0.0001, Figure 2Civ). Similarly, MNNG-HOS+P2RX7B cells showed a 15% increase in wound closure when stimulated with 10 µM BzATP (P= 0.002, Figure 2Cii and 2Cvi). To measure the invasion ability of OS cells a Fluroblok transwell system containing matrigel was used with 10% FBS in the lower chamber as a chemoattractant and the cells in the upper chamber stimulated with 10µM BzATP (as this induced increased migration in both cell lines). The TE85 cell line is a relatively non-invasive cell line therefore only a few detectable cells invaded the transwell (Figure 2Di). Transfection with P2RX7B and stimulation with BzATP provided a strong stimulus to promote invasion increasing this from an average of 49 cells to 298 cells (P=0.00374, Figure 2Dii). MNNG-HOS cells showed a higher level of invasion in keeping with their phenotype (Figure 2Diii), with a similar increased invasion seen in the MNNG-HOS+P2RX7B cells (increasing from an average of 278 cells to 662 cells, P=0.025, Figure 2Div).

### Effect of P2RX7B expression and inhibition on tumour growth and bone disease in an OS mouse model *in vivo*

To determine whether P2RX7B expression would affect the growth and progression of OS disease *in vivo* a murine paratibial model of OS was used. This model has the advantage of an intact bone at the outset and results in tumours that reproduce the most common osteoblastic form of human OS^39-41^. As TE85 cells do not form tumours *in vivo*, and this remained the case even after transfection with P2RX7B (data not shown), we concentrated the *in vivo* part of this study using the MMNG-HOS cell lines. Injection of either MNNG-HOS or MNNG-HOS+P2RX7B cells resulted in the formation of OS in the tibia (Figure 3A). There was no statistical difference in the final tumour size between mice injected with MMNG-HOS+P2RX7B cells compared to MMNG-HOS cells (P=0.3071, Supplementary Figure 2Ai). The P2RX7 antagonist A740003 did reduce the size of the tumours from both cell lines although this did not quite reach statistical significance (P=0.0568 for MNNG-HOS+P2RX7B and P=0.0516 for MNNG-HOS; Supplementary Figure 2Aii and 2Aiii). Ki-67 staining of the tumours similarly demonstrated no significant difference in the percentage of cells stained positive in MNNG-HOS+P2RX7B tumours compared to MNNG-HOS tumours (P=0.6329 Supplementary Figure 2Bi) or when either group was treated with A740003 (P=0.9074 Supplementary Figure 2Ci and P=0.4010 respectively, Supplementary Figure 2Di). When divided into high and low grade tumours, there was no difference in the percentage of grades between the cell lines (P=0.4201 Supplementary Figure 3Bii).

**Figure 3:**
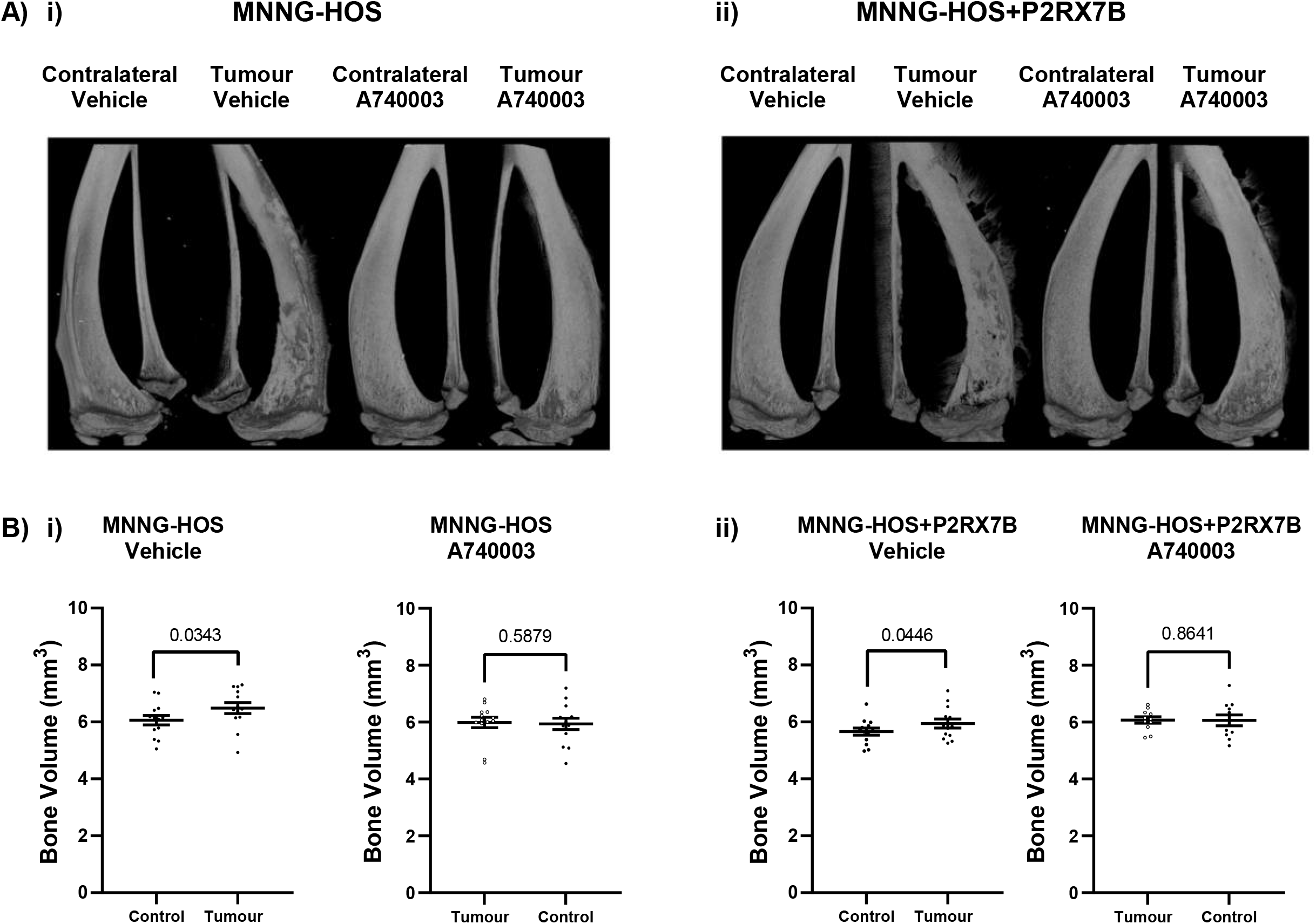
P2RX7B expression and P2RX7B inhibition on bone disease in an OS mouse model *in vivo*. Female BALB/c 7-9 week old mice were injected paratibially with either MNNG-HOS or MNNG-HOS+P2RX7B cells (250,000 cells suspended in 20 µL PBS). Mice were then randomised into groups and treated with either vehicle or A740003 from day 2, three times a week for 3 weeks via IP injection. Calliper measurements were taken on the final day as an end-point tumour measurement prior to euthanasia, the legs were then collected and micro-CT scanned, the total bone volume starting from the point at which the femur wasn’t visible and the fibula meets the tibia was determined for each leg and normalised to its own contralateral leg. Quantification of the bone volume (mm^3^) was done for all groups. n= 11-13 per group, the difference between the tumour bearing and contralateral control leg determined by a paired t-test appropriate for the distribution of the data in the groups, and a Two-way ANOVA was used to determine the effect of P2RX7B expression on tumour-induced changes in bone parameters. **Ai)** Representative micro-CT images of MNNG-HOS tumour bearing and contralateral leg receiving vehicle and A740003 with the corresponding bone volumes shown in **Bi**. Representative micro-CT images of MNNG-HOS+P2RX7B tumour bearing and contralateral leg receiving vehicle and A740003 are shown in **Aii)** and the corresponding bone volumes shown in **Bii**.

When both cells lines were treated with the P2RX7 antagonist A740003, the ratio of low to high grade tumours improved in both cell lines, however the association was not statistically significant, (P=0.1156 for MNNG-HOS and P=0.3917 for MNNG-HOS+P2RX7B respectively Supplementary Figure 2Cii & 3Dii).

A defining feature of OS is the involvement of the bone itself, therefore resultant effect on bone was measured using micro-CT. Injection of OS cells led to the typical ectopic bone formation (Figure 3A) with a significant increase in total bone volume of the injected leg compared to its contralateral leg for both MNNG-HOS (P=0.0343, Figure 3Bi) and MNNG-HOS+P2RX7B cells (P= 0.0446, Figure 3Bii). When considering the effect of the cell type of OS tumour on the bone disease, Two-Way ANOVA revealed that whilst the OS tumour accounted for 7.758% of the variance (P=0.0341), expression of P2RX7B in the MNNG-HOS cells accounted for 13.62% of the variance (P=0.0058). Treatment with A740003 reduced the extent of the tumour induced bone disease both in MNNG-HOS and MNNG-HOS+P2RX7B such that the bone volume was no longer significantly increased compared to the contralateral leg (P=0.2939, and P=0.4783 respectively, Figure 3B).

### P2RX7B expression increases spontaneous OS lung metastasis *in vivo*

To investigate the role of P2RX7B in OS metastasis the lungs were collected from each mouse from the paratibial model and examined histologically for signs of micrometastasis. We did not detect metastasis in any of the 26 mice bearing MNNG-HOS primary OS tumours, but we did detect metastasis in mice bearing MNNG-HOS+P2RX7B primary OS tumours. Out of 24 mice with primary OS tumours, 5 (21%) had incidence of metastasis representing a significant difference (p=0.0142) in the incidence of detectable lung metastasis between the two cell lines (Figure 4A & 4E). P2RX7 antagonist treatment slightly reduced the incidence of metastasis from 3/13 (23%) mice for vehicle to 2/11 (18%) mice for A740003 (Figure 4B & 4E) but this did not reach statistical significance.

**Figure 4:**
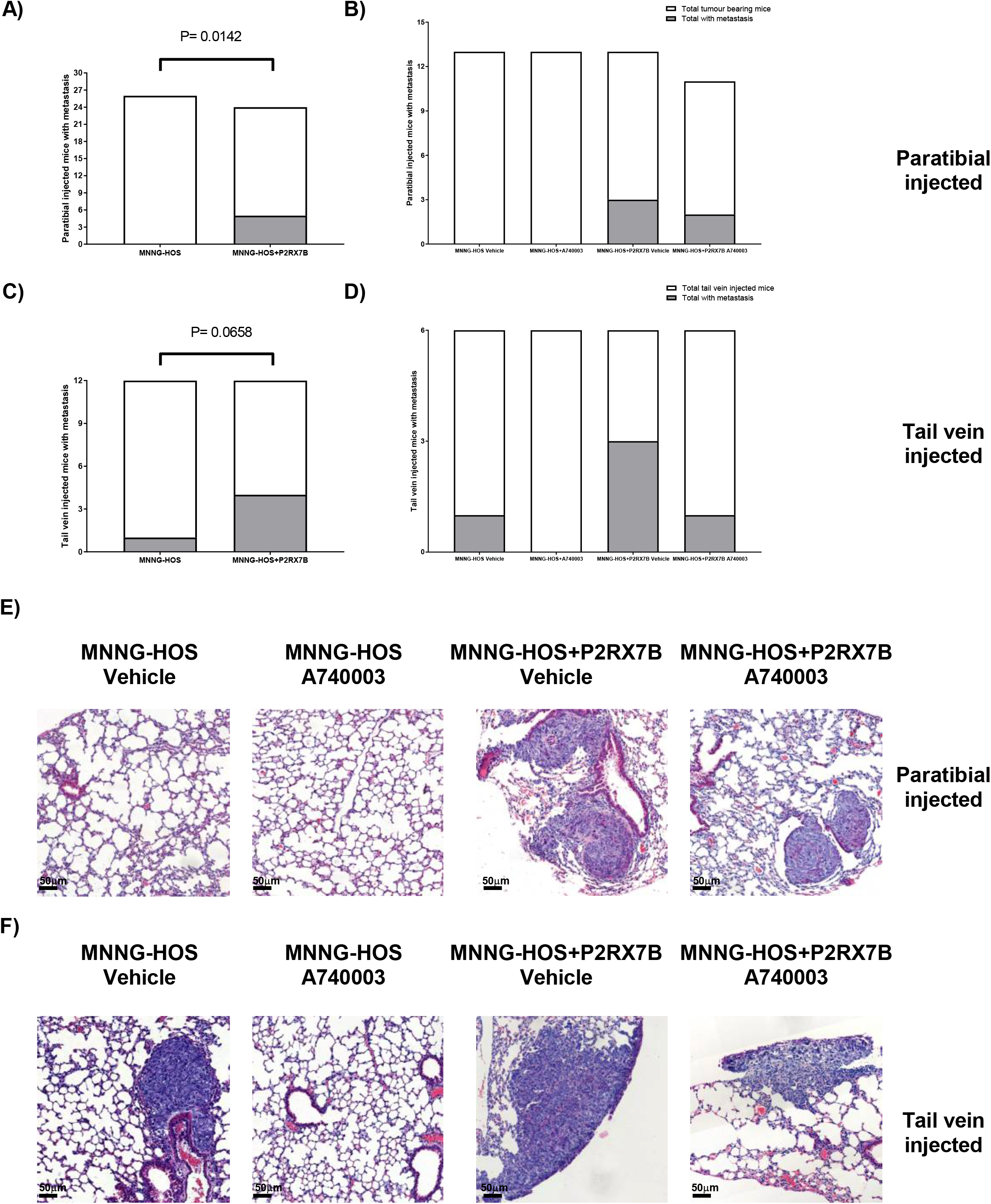
P2RX7B expression increases OS metastasis in the lungs of MNNG-HOS and MNNG-HOS+P2RX7B paratibial and tail vein injected mice. Female BALB/c 7-9 week old mice were injected paratibially with 250,000 cells suspended in 20 µL PBS of either MNNG-HOS or MNNG-HOS+P2RX7B cells. Mice were then randomised into groups and treated with either vehicle or A740003 from day 2, three times a week for 3 weeks via IP injection. For the tail vein model the same treatment regime was used after 1.5 million cells were injected. The mice were then euthanised and the lungs embedded into wax blocks sectioned and 6 μm sections cut every 100 μm deep to cover the entire lung. They were then stained using H&E and visualised using a light microscope. **A**) Total number of OS lung metastasis in MNNG-HOS and MNNG-HOS+P2RX7B tumour bearing mice in the partibial model. **B)** Number of OS lung metastasis in MNNG-HOS and MNNG-HOS+P2RX7B tumour bearing mice across treatment groups in the partibial model. **C)** Total number of OS lung metastasis in MNNG-HOS and MNNG-HOS+P2RX7B injected mice in the tail vein model. **D)** Number of OS lung metastasis in MNNG-HOS and MNNG-HOS+P2RX7B injected mice across treatment groups in the tail vein model. **E)** Representative images of the lungs from MNNG-HOS and MNNG-HOS+P2RX7B vehicle and A740003 treated in the paratibial model **F)** from MNNG-HOS and MNNG-HOS+P2RX7B vehicle and A740003 treated in the tail vein model.

To further explore the role that P2RX7B plays in OS metastasis an experimental metastasis study was performed where mice were tail vein injected with either MNNG-HOS or MNNG-HOS+P2RX7B cells and treated with either vehicle or A740003. The incidence of lung metastasis using this model was again higher in the mice injected with MNNG-HOS+P2RX7B cells (4/12 or 33%) compared to the mice injected with MNNG-HOS (1/12 or 8%) (Figure 4C & 4F, p=0.0658). Treatment with the P2RX7 antagonist treatment reduced the incidence of metastasis to none in the MNNG-HOS cell injected mice, and from 3/6 (50%) to 1/6 (∼17%) for MNNNG-HOS+P2RX7B injected mice (Figure 4D & 4F), although this did not reach statistical significance.

### P2RX7B inhibition drives downregulation of *FN1*/*LOX*/*PDGFB*/*IGFBP3*/*BMP4* axis

To determine the downstream pathways contributing to the observed effects of P2RX7B signalling in OS cells *in vitro* and *in vivo*, we took a hypothesis free approach using RNA-seq. We performed heat map based hierarchical cluster analysis on DE genes unique to MNNG-HOS+P2RX7B cells when either treated with BzATP (Figure 5A) or the antagonist A740003 (Figure 5B). These genes were cross-referenced with several control cell lines to ensure that our data was derived from P2RX7B activation/inhibition and from not off target effects (Supplementary File). P2RX7B activation and antagonism were shown to drive independent responses, with antagonism specifically demonstrating a more potent biological response when in the context of cancer, showing a visibly higher number of genes affected and these are strongly related to cancer progression (Figure 5 A-B). STRING analysis of gene–gene connections at high confidence level revealed an axis of downregulated genes in the P2RX7B antagonist treated MNNG-HOS+P2RX7B cells (Figure 5B). This analysis revealed that P2RX7B antagonism downregulated an *FN1*/*LOX*/*PDGFB*/*IGFBP3*/*BMP4* axis, which contains genes that are all known cancer drivers when upregulated^42^.

**Figure 5:**
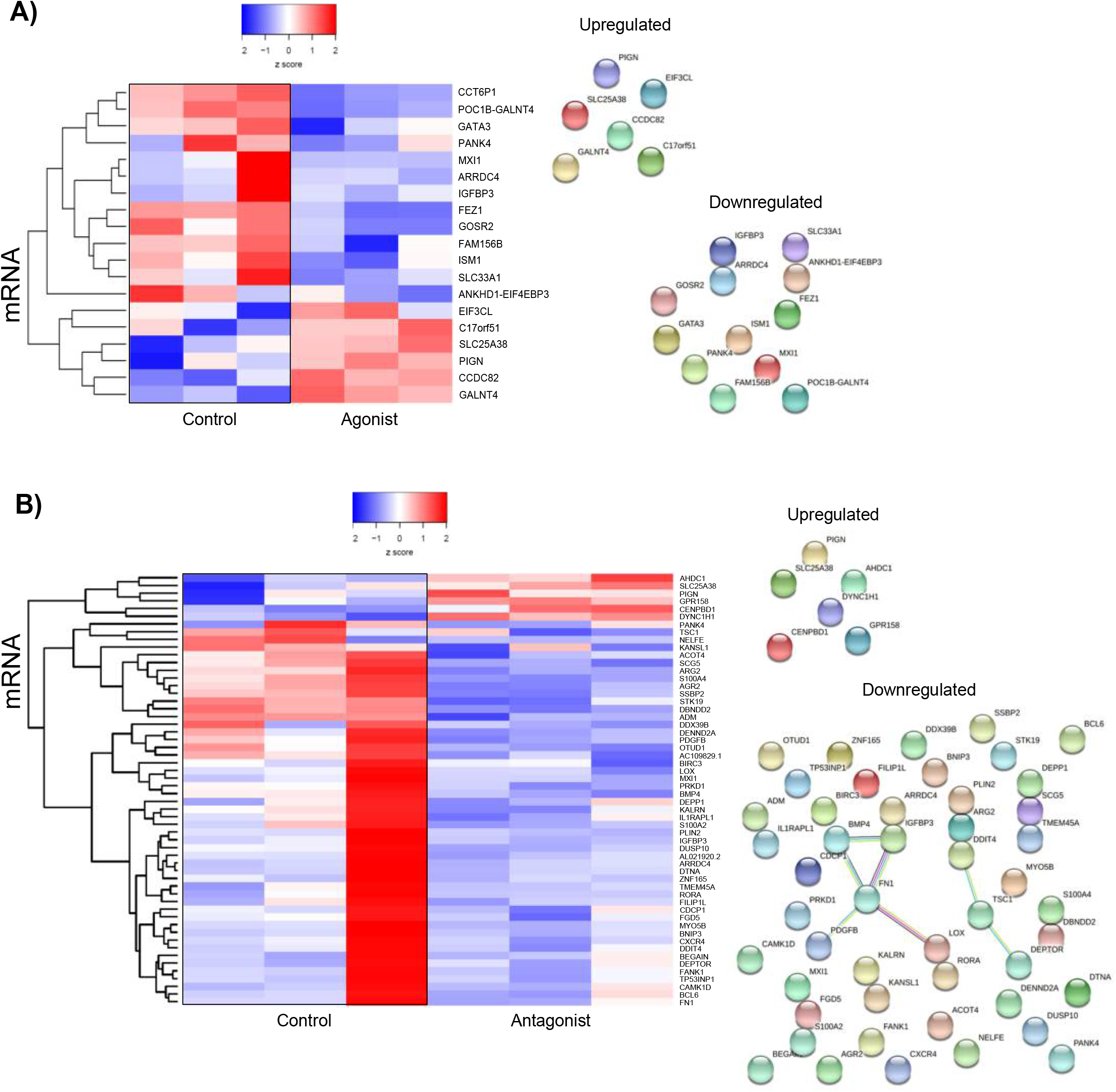
Heat map based hierarchical cluster analysis of DE genes (y-axis) across cell type (y-axis). Z-score refers to high (red) and low (blue) gene expression using normalised values when compared to the mean of total sequencing reads. Heat maps are accompanied by STRING analysis to show gene–gene connections at high confidence (scores between 0.7 and 0.9). Line colour connecting genes indicate the known and predicted interactions. Blue lines represent data from curated databases. Pink lines represent data from experiments. Green lines represent gene neighbourhoods. Black lines represent co-expressed genes. **A)** MNNG-HOS+P2RX7B cells versus activated receptors using BzATP. **B)** MNNG-HOS+P2RX7B cells versus inhibited receptor using A740003 antagonist. Potential off target effects were considered by performing RNA-seq in MNNG-HOS cells (Supplementary File). Each transcript presented has passed log_2_ fold change ≥2, *p* ≤0.05 and FDR ≤ 5% parameters.

## Discussion

Despite being the most common type of primary bone cancer affecting children and adolescents, treatment options for OS have remained stagnant for the last 40 years or more. The overall five-year survival for OS is 60% largely due to the fact that many patients relapse from chemoresistance or have metastasis on diagnosis reducing the survival rate in these patients to around 20%^3^. Treatment options are brutal and life changing, with surgery often leading to permanent disability and chemotherapy having significant side effects including infertility, cardio-, nephro- and hepatic-toxicity. More targeted treatment options are needed to effectively treat OS patients with fewer treatment side effects.

We and others have previously reported the role of P2RX7 purinergic signalling in both the bone microenvironment and OS^8, 9, 15, 35^, and this particular receptor has been increasingly implicated in cancer progression in various cancer types^43^. In this study we have built upon these observations and detailed the role of the P2RX7B splice variant specifically in OS. Having previously demonstrated that the cell line TE85 does not express functional P2RX7^15, 35^ we chose to use this cell line as well as the sub-clone MNNG-HOS to investigate the effect of the splice variant P2RX7B alone on OS cell behaviour. Prior to transfecting these cells with the P2RX7B, we confirmed that neither cell line expressed P2RX7 at the mRNA level. This finding is in direct contrast to a recent study, published during our own ongoing studies, which suggested that MNNG-HOS cells highly express P2RX7^24^. We also scrutinised publicly available datasets to determine the relative levels of P2RX7 in the HOS family of OS cell lines and found that HOS, MNNG-HOS, KHOS and 143B cells all had low or non-existent expression of P2RX7, further confirming our observations.

Following transfection of P2RX7B into TE85 and MNNG-HOS cells, end point PCR and qRT-PCR confirmed robust mRNA expression. There was also an increased calcium response when stimulated with BzATP in P2RX7B transfected clones. We also checked BzATP-induced pore formation in these cells, as despite P2RX7B not having the typical pore formation response due to a truncated C terminal region known to be responsible for the pore, we have previously demonstrated that co-expression of P2RX7A and P2RX7B does lead to pore formation^15^. This further demonstrates a lack of endogenous P2RX7A expression in these cells, that in our clones the expression increase relates to P2RX7B and in these clones a “ non-functional” phenotype is present^20^.

Having established our P2RX7B expressing OS cell lines, we evaluated the effect of receptor expression on typical cancer cell behaviours. As with our previously published observations in TE85 cells^15^, we confirmed a strong growth advantage when P2RX7B was expressed in both TE85 and also MNNG-HOS cells. This growth advantage was observed in low serum conditions without exogenous activation of the receptor – suggesting an autonomous activation of P2RX7B due to basal release of ATP and autocrine/paracrine signalling known to occur in cells, including bone and cancer cells^44^. The finding that the P2RX7 antagonists A740003 could significantly reduce this growth advantage offers promise for inhibiting P2RX7B mediated effects.

We next sought to determine the effect of P2RX7B expression on the ability of OS cells to leave the primary tumour site, invade surrounding tissues and spread to distant sites to form metastasis – the so called “ invasion-metastasis cascade” ^45^. This is particularly pertinent in OS and other highly metastatic cancers, as the 5-year survival rate is drastically reduced in patients with metastasis^3^. Using established *in vitro* techniques, we found that P2RX7B expression reduced the ability of OS cells to adhere to a collagen ECM. Non-cancerous cells are normally tightly adherent to their ECM and also to each other – changes in cell-to-cell and cell-to-matrix adhesion is the first fundamental step in the metastatic cascade. E-cadherin is known to be pivotal in the control of cancer cell adhesion and spread, and it has previously been demonstrated that PX7R activation downregulates E-cadherin in breast cancer cells^46^ Loss of E-cadherin mediated adhesion is also frequently associated with invasion and migration. Similar mechanisms involving cadherin signalling could be responsible for the P2RX7B-induced loss of adhesion in OS cells. The P2RX7B expressing OS cells when stimulated with the potent P2RX7 agonist BzATP, demonstrated increased invasion and migration rates. Invasion requires the expression of various matrix metalloproteinases (MMPs) that are capable of digesting the ECM, allowing the cells to break away from the primary site and interact with tumour-associated stromal cells such as macrophages^47^. Activation of P2RX7 has been shown to induce the release of MMP13 in breast cancer cells^46^ and has long been known to be responsible for the ATP-induced rapid release of MMP-9 from human peripheral-blood mononuclear cells^48^. This data further supports a pro-tumorigenic role of P2RX7 in OS as demonstrated previously^15, 24^, but adds new knowledge that the B variant alone plays a role in the metastasis cascade in OS cells *in vitro*.

Encouraged by the *in vitro* data, we next examined the role of P2RX7B in OS *in vivo*. As previously mentioned, the TE85 cell line does not produce tumours in mice therefore the MNNG-HOS cell lines only were used. Assessment of the growth of the primary tumour using caliper measurements and Ki-67 staining did not show any effect of P2RX7B expression or any effect of the P2RX7 antagonist A740003 on the final tumour volume or proliferation status. This was a surprising finding given the *in vitro* data and the previous study from Zhang *et al*. who showed a 50% reduction in the growth of intra-tibial MNNG-HOS tumours with A740003 treatment^24^. The differences between our data and theirs maybe explained by the different OS models used (paratibial versus intra-tibial injection of cells), timing of treatment (2 days post cell injection versus ∼10 days) and also potential differences in the parental MNNG-HOS cell lines. As stated above we could not find any P2RX7 mRNA expression or typical function in the MNNG-HOS cells we used. When analysing the resultant bone disease in our OS model, expression of P2RX7B significantly affected the bone disease observed, with A740003 reduced the extent of the tumour induced bone disease in both MNNG-HOS+P2RX7B and MNNG-HOS such that the bone volume was no longer significantly increased compared to the contralateral leg. Whilst this is in keeping with the positive effect of P2RX7 antagonism on OS-induced bone disease previously reported, it is important to note again the difference between the two models. Here we show for the first time the specific involvement of P2RX7B in OS induced bone disease; that MNNG-HOS+P2RX7B cells produce an osteoblastic phenotype with formation of ectopic bone and so an increase in total bone volume which was reduced by treatment with A740003. Zhang *et al* show that the MNNG-HOS cells they used produced predominantly a lytic bone disease leading to reduced BV/TV which was also rescued with A740003 treatment^24^. We also observed a reduction in the ectopic bone formation in the parental MNNG-HOS cells when treated with A740003. Considering that in our hands these cells do not express endogenous, functional P2RX7 receptors and the lack of effect on tumour size, the effect of A740003 may be mediated by the bone microenvironment itself, as opposed to a direct tumour driven effect. We and others have shown that both osteoblasts and osteoclasts express functional P2RX7 receptors^8^, and the P2RX7 knockout mice display a bone phenotype^49-52^. Therefore treatment with A740003 could be affecting the resident bone cells and reducing the resultant bone disease in both subtypes of OS. Targeting the bone microenvironment in OS is a valid strategy, with other mediators of osteoblast and osteoclasts being previously proposed as potential therapeutic targets^53^. The five year survival rate for OS in the presence of secondary metastases is around 20%. In addition, a quarter of patients presenting with detectable lung metastasis relapse. Gaining further insight into the potential mechanisms or predictors of OS metastasis is of paramount importance. By examining the lungs of the tumour bearing mice we found that P2RX7B expression in the primary tumour led to a significantly higher incidence of lung metastasis. This is in contrast to the lungs of the parental MNNG-HOS tumour bearing mice which showed no metastasis, in keeping with the previous observations that MNNG-HOS cells *in vivo* have no or very low incidence of pulmonary metastasis^36, 38^. Taken together with the *in vitro* data of decreased adhesion with increased invasion and migration abilities of the MNNG-HOS+P2RX7B cells and the minimal effect on tumour growth seen in our *in vivo* studies, these data support a role for the P2RX7B in the metastasis cascade in OS.

Finally, to try and elucidate the pathways downstream of P2RX7B in OS we utilized a hypothesis-free RNA-seq approach. Using this methodology we revealed a set of genes that are differentially expressed upon both P2RX7B activation and antagonism. Antagonism of P2RX7B showed a more potent response in the context of cancer leading to a higher number of genes affected, but also suggesting that P2RX7B has activation state-dependent roles. These genes were strongly related to cancer and STRING analysis revealed a unique axis of downregulated genes in the antagonist treated MNNG-HOS+P2RX7B cells. Within this axis the genes *FN1, LOX, PDGFB, IGFBP3* and *BMP4* are known to promote cancer progression when upregulated in various cancer types. Roles for these genes/gene products in OS have also been reported: *FN1* was shown to be upregulated in chemoresistant OS cells, was related to unfavourable prognosis and its inhibition greatly increased the sensitivity of OS cells to doxorubicin *in vitro* and *in vivo*^54^. LOX (Lysyl Oxidase) has been suggested to be tumour suppressor in OS^55^, however we and others have shown that it is linked to hypoxia and plays a pro-tumorigenic role in cancers that metastasis to bone, inducing the pre-metastatic niche both in soft tissues such as lungs and in bone via formation of osteolytic lesion^56^. PDGFB is also linked to hypoxia and has been shown to be involved in OS cell proliferation and migration^57^, and *IGFBP3* has also been shown to stimulate OS migration^58^. The *BMP4* protein, part of the bone morphogenetic family which are known critical regulators of bone formation and remodelling, has recently been shown to be the most frequently expressed BMP in musculoskeletal tumours, including OS, in a recent study of 1,361 patients with 22 types of musculoskeletal tumours^59^. Elucidating the interplay between P2RX7B and these signalling pathways could reveal common targets at which they converge and that may prove to be fundamental in OS progression and metastasis. Further studies should look to determine the effect of combination treatments (both within this axis and with current OS drugs) that may prove to be more efficacious than single agent treatments. Combining treatments for signalling pathways specifically involved in OS would lead to kinder, less harsh treatments for the patient, fewer relapses and a better overall prognosis.

In summary we present data for the first time that the specific P2RX7B isoform, which has a distinct non-functional phenotype compared with full length P2RX7, is associated with OS progression, bone disease and metastasis. Future studies exploring the therapeutic targeting of P2RX7 need to pay particular attention to the isoforms being targeted as well as the OS subtype.

## Acknowledgements

The authors would like to thank the Bone Cancer Research Trust for supporting this work, grant BCRT Grant 4415 awarded to A.G.

## Abbreviations (alphabetical)

ATP: Adenosine Triphosphate
BzATP: Benzoyl ATP
DE: Differentially Expressed
ECM: Extracellular Matrix
FDR: False Discovery Rate
HBSS: Hank’s Balanced Salt Solution
LOX: Lysyl Oxidase
MMPs: Matrix Metalloproteinases
OS: Osteosarcoma
PE: Paired End
qRT-PCR: Quantitative Reverse Transcription Polymerase Chain Reaction

**Supplementary 1: P2RX7 expression in different OS cell lines determined through In silico analysis of publically available datasets**.

A retrospective analysis across publically available datasets was performed to identify P2RX7 expression in human OS cell lines. Searches were performed in: **A)** Ordino a visual cancer analysis database. **B)** The Cancer Dependency Map 20Q4 depmap gene expression dataset. **C-F)** R2 Genomics Analysis and Visualization Platform, **C)** Data set GSE42352 **D)** Data set GSE124768 **E)** Data set GSE86109 **F)** Data set GSE11414. The gene P2RX7 was selected and filtered for OS cell lines then selected and included. Cell lines relating to the HOS lineage are highlighted in red.

**Supplementary Figure 2: P2RX7B expression and P2RX7B inhibition on tumour volume, Ki-67 staining and histological high/low grading of tumour sections in an OS mouse model *in vivo***

Female BALB/c 7-9 week old mice were injected paratibially with either MNNG-HOS or MNNG-HOS+P2RX7B cells (250,000 cells suspended in 20 µL PBS). Mice were then randomised into groups and treated with either vehicle or A740003 from day 2, three times a week for 3 weeks via IP injection. At sacrifice the tumour size was measured and the legs collected, processed, embedded into wax blocks and sections were taken to be stained for Ki-67. The sections were then analysed using QuPath to determine percentage Ki-67 staining and divided into high or low grade tumours based on their median value N= 11-13 per group. **Ai)** MNNG-HOS tumour size compared to MNNG-HOS+P2RX7B **Aii)** MNNG-HOS+P2RX7B mice treated with A740003 **Aii)** MNNG-HOS mice treated with A740003. **Bi)** Ki-67 percentage for MNNG-HOS mice compared to MNNG-HOS+P2RX7B with histologically graded tumours compared in **Bii. Ci)** Ki-67 percentage for MNNG-HOS mice treated with vehicle and A740003, with histologically graded tumours in **Cii. Di)** Ki-67 percentage for MNNG-HOS+P2RX7B mice treated with vehicle and A740003 with histologically graded tumours in **Dii**. Ki67 percentage results were compared using an unpaired T-Test and Chi-Squared was used for histologically graded tumours with representative images for MNNG-HOS tumour bearing mice shown in **Ciii** and MNNG-HOS+P2RX7B tumour bearing mice shown

