## Supplementary figures and images for "The P2RX7B splice variant modulates osteosarcoma cell behaviour and metastatic properties"

### Supplementary 1

#### Supplementary 1

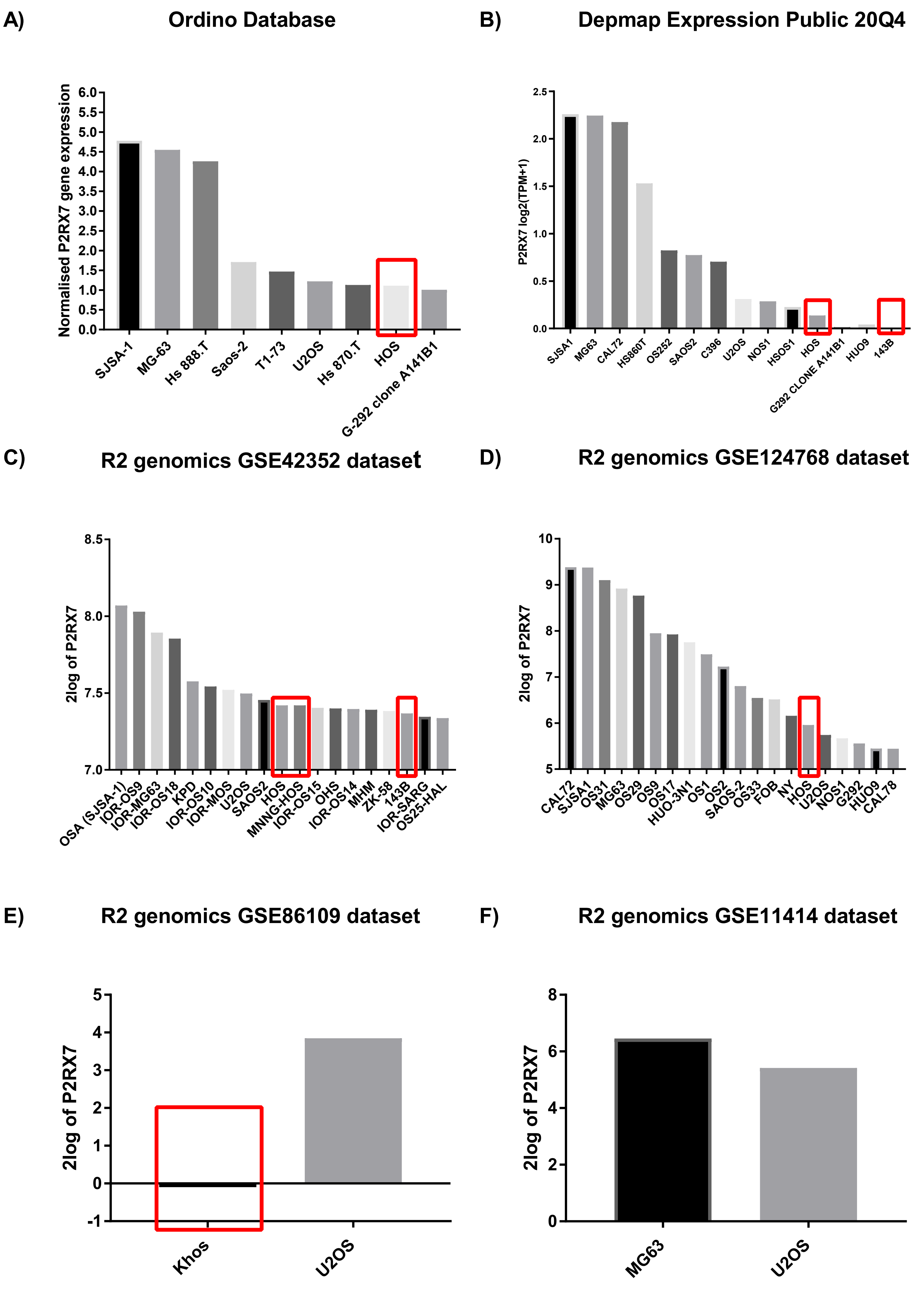

### Supplementary 2

#### Supplementary 2

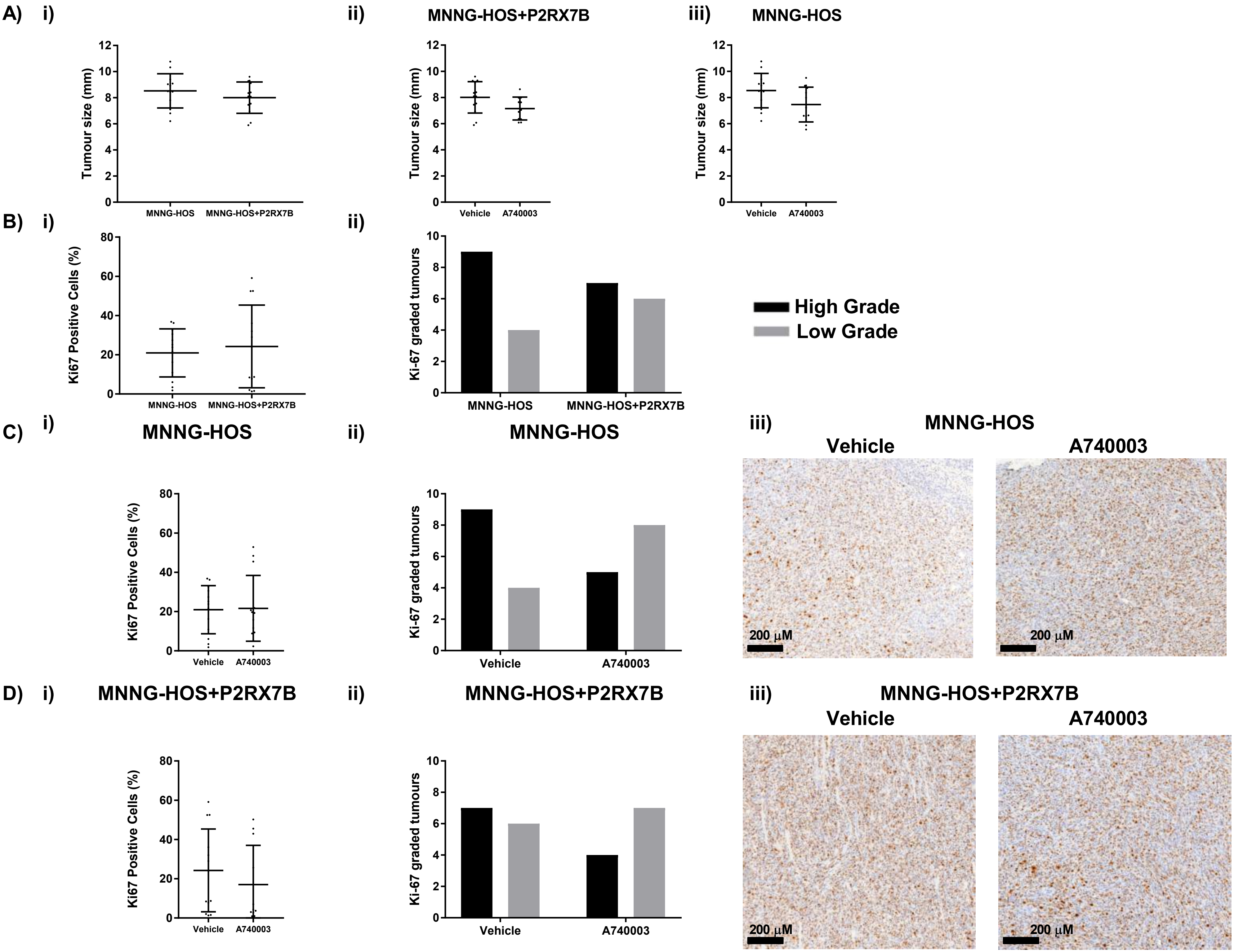
